# γδ T cell-derived IL-17A aggravates experimental epidermolysis bullosa acquisita in response to IL-1β

**DOI:** 10.64898/2026.11.04.763966

**Authors:** Fengyuan Deng, Sandra Jukic, Melanie Wannick, Alexandra Hölscher, Immo Prinz, Ralf J. Ludwig, Enno Schmidt, Xinhua Yu, Frank Petersen, Christoph Hölscher

**Affiliations:** Infection Immunology, Research Center Borstel, Borstel, Germany; Key Laboratory of Basic Research on Regional Diseases, Guangxi Medical University, Guangxi, China; Institute of Systems Immunology, Hamburg University Medical Center, Hamburg-Eppendorf, Germany; Institute of Experimental Dermatology, University of Lübeck, Lübeck, Germany; Pulmonary Immune Diseases, Research Center Borstel, Borstel, Germany

**Keywords:** epidermolysis bullosa acquisita, interleukin-17A, interleukin-1β, gamma delta T cells, antibody-transfer model

## Abstract

Epidermolysis bullosa acquisita (EBA) is a mucocutaneous autoimmune blistering disorder mediated by autoantibodies against type VII collagen and subsequent splitting of the dermal-epidermal junction. Interleukin (IL)-17A promotes the production of chemokines and cytokines involved in neutrophil recruitment and activation, suggesting that IL-17A may contribute to EBA pathogenesis. We aimed to clarify the role of IL-17A in experimental EBA. Using an antibody transfer-induced EBA mouse model, we demonstrate that IL-17A promotes disease development, as pharmacological blockade of IL-17A with a neutralizing antibody ameliorated skin lesions. Gamma-delta (γδ) T cells were identified as the predominant cellular source of IL-17A, supported by the attenuated disease phenotype observed in γδTCR-deficient mice. In addition, CD4^+^ T cells and neutrophils contributed to IL-17A production in experimental EBA. Mechanistically, IL-1-beta, rather than IL-23, was identified as a critical mediator of IL-17A production by γδ T cells *in vitro*, consistent with our *in vivo* findings. Whereas EBA severity was not reduced in IL-23p19-deficient mice, disease was markedly attenuated in IL-1R-deficient animals. Together, our study provides the first evidence that γδ T cell-derived IL-17A plays a pivotal role in experimental EBA and identifies a previously unrecognized IL-1β/γδ T cell/IL-17A axis that expands current understanding of EBA immunopathogenesis.

## INTRODUCTION

Epidermolysis bullosa acquisita (EBA) is a chronic autoimmune bullous disease caused by autoantibodies against type VII collagen (COL7), the major structural component of anchoring fibrils at the dermal–epidermal junction (Koga et al., 2018; Woodley et al., 1984). These autoantibodies compromise the structural integrity of the basement membrane zone, resulting in skin fragility, blister formation, and chronic wounds that markedly impair quality of life (Ludwig and Zillikens, 2011). Histologically, affected skin from patients is characterized by prominent granulocyte infiltration and deposition of IgG and complement (Kim and Kim, 2013). The pathogenic cascade in EBA is initiated by binding of pathogenic autoantibodies to COL7 at the dermal–epidermal junction, followed by complement activation, release of cytokines and chemokines, and recruitment of inflammatory cells such as T cells, macrophages, and particularly granulocytes (Ludwig and Zillikens, 2011).

To elucidate the pathogenesis of EBA, several mouse models have been established, either by immunization with recombinant murine COL7 (mCOL7) or by passive transfer of antibodies against mCOL7 (Sitaru et al., 2006; Sitaru et al., 2005; Woodley et al., 2005; Woodley et al., 2006). These experimental models have substantially advanced our understanding of how autoantibodies against COL7 induce inflammation and tissue damage. Several cellular and molecular components have been shown to contribute to experimental EBA. The production of anti-COL7 autoantibodies is driven by a CD4^+^ T cell-dependent B-cell response, whereas their persistence is regulated by the neonatal Fc receptor (Iwata et al., 2013; Sesarman et al., 2008; Sitaru et al., 2010). Following autoantibody binding to COL7, tissue injury and blister formation are mediated by multiple pathogenic mechanisms, including complement activation, particularly through C3a and C5a, the production of pro-inflammatory cytokines such as interleukin (IL)-6 and IL-8, neutrophil recruitment and activation, and the release of reactive oxygen species and proteases (Hirose et al., 2013; Mihai et al., 2007; Sadeghi et al., 2015; Samavedam et al., 2014; Syed et al., 2009; Yu et al., 2018). Neutrophils are widely recognized as essential effector cells mediating tissue damage in EBA pathogenesis, as neutrophil-depleted mice fail to develop blistering after passive transfer of anti-COL7 antibodies (Chiriac et al., 2007).

IL-17A is a multifunctional pro-inflammatory cytokine produced by multiple immune cell types, including CD4^+^ T-cell subsets, gamma-delta (γδ) T cells, CD8^+^ T cells (Tc17 cells), natural killer T cells, and innate immune cells such as neutrophils and monocytes (Mills, 2023). Through its heterodimeric receptor complex composed of IL-17 receptor (R)A and IL-17RC, IL-17A activates downstream NF-κB and mitogen-activated protein kinase signaling pathways, thereby inducing neutrophil chemoattractants, including C-X-C motif chemokine ligand (CXCL)1, CXCL2, and CXCL8/IL-8, pro-inflammatory cytokines such as IL-6, IL-1 beta (β), and Tumor-Necrosis-Factor (TNF), and matrix-degrading proteases (Veldhoen, 2017; Xu Lou et al., 2025). IL-17A is therefore considered a potent orchestrator of neutrophil-dominated inflammation: it promotes myelopoiesis via induction of granulocyte colony-stimulating factor (G-CSF) and granulocyte-macrophage colony-stimulating factor (GM-CSF) while simultaneously stimulating keratinocytes to secrete neutrophil-attracting chemokines (Ye et al., 2001). Consistently, increased IL-17A expression has been reported in several neutrophil-associated autoimmune skin diseases, including psoriasis, bullous pemphigoid (BP), and pemphigus vulgaris (Chakievska et al., 2019; Kutwin et al., 2021; Song and Qian, 2013; Xue et al., 2014; Zhu and Qian, 2012).

The critical involvement of IL-17A in neutrophilic inflammation has been firmly established in psoriasis, where IL-17A-targeted biologics, including secukinumab, ixekizumab, and brodalumab, have substantial (Navarro-Compán et al., 2023)ly improved therapeutic outcomes (Brembilla et al., 2018). In BP, the most common autoimmune blistering disease, elevated IL-17A levels have been detected in patient sera, blister fluid, and lesional skin (Arakawa et al., 2011; Chakievska et al., 2019). Moreover, Chakievska and colleagues demonstrated that IL-17A-deficient (^-/-^) mice are protected from experimental BP and that pharmacological inhibition of IL-17A significantly reduces disease severity (Chakievska et al., 2019). In individual BP patients and small cohorts, IL-17 inhitionhas shown promising clinical effects (Grochowska-Rak et al., 2026; Heidari et al., 2025; Holtsche et al., 2020). These findings suggest that IL-17A is a key mediator of skin and mucosal immune responses and contributes substantially to neutrophil-mediated inflammation.

Although EBA is a prototypic inflammatory blistering disorder, the functional contribution of IL-17A to EBA remains insufficiently defined. We therefore hypothesized that IL-17A promotes the development of experimental EBA.

To test this hypothesis, we used the antibody transfer model of EBA to determine the relative contribution of IL-17A, identify its cellular source, and define upstream mechanisms regulating IL-17A production during antibody-induced EBA.

## RESULTS

### Pharmacological inhibition of IL-17A significantly reduces skin inflammation and blistering in an antibody transfer-induced mouse model of EBA

Experimental EBA was induced in C57BL/6 mice by repeated injections of anti-mCOL7 IgG or normal rabbit serum on days 0, 2, 4, 6, 8, and 10. To block IL-17A function, mice were treated with an anti-IL-17A antibody or an isotype control antibody on days −2, 0, 2, 4, 6, 8, and 10. EBA manifestations, including erosions and blister formation, were observed in both anti-mCOL7 IgG-treated groups; however, compared with mice receiving the isotype control antibody, mice treated with anti-IL-17A antibody showed markedly ameliorated disease (Figure 1A). Disease severity was evaluated as the percentage of affected body surface area on days 4, 8, and 12 after the initial injection. On day 12, anti-IL-17A-treated mice displayed a significantly lower percentage of affected body surface area than control mice (4.919 ± 3.141% *vs* 1.398 ± 0.732%, Figure 1B, p < 0.05). Histopathological changes in skin lesions were analyzed by H&E staining of formalin-fixed ear skin sections, and neutrophil infiltration into affected skin was assessed by immunohistochemistry (Figure 1A). In isotype-treated mice receiving anti-mCOL7 IgG, H&E staining revealed pronounced epidermal disruption with dense dermal inflammatory infiltrates, accompanied by prominent neutrophil accumulation in lesional skin. In contrast, IL-17A blockade markedly reduced histopathological tissue damage and neutrophil infiltration, consistent with the attenuated clinical phenotype. To confirm autoantibody binding and complement deposition at the dermal–epidermal junction, immunofluorescence staining was performed on skin cryosections collected on day 12. IgG binding and C3 deposition were observed in anti-mCOL7 IgG-injected groups, but not in normal rabbit IgG-injected groups (Figure 1C). These findings indicate that IL-17A blockade attenuates inflammatory tissue injury without impairing autoantibody binding or local complement deposition. To assess whether the protective effect of IL-17A blockade was accompanied by changes in local inflammatory mediator expression, transcript levels of selected cytokines and chemokines were analyzed. mRNA levels of *Il6* and *Ccl2* showed a tendency toward reduction in IL-17A-blocked mice compared with control C57BL/6 mice after anti-mCOL7 IgG injection; however, these differences did not reach statistical significance. In contrast, *Cxcl1* expression was comparable between both groups (Figure 1D). Together, these findings demonstrate that pharmacological inhibition of IL-17A markedly attenuates clinical disease severity and inflammatory tissue damage in antibody transfer-induced EBA, while autoantibody binding, complement deposition, and the expression of selected inflammatory mediators remain largely unaffected.

**Figure 1.**
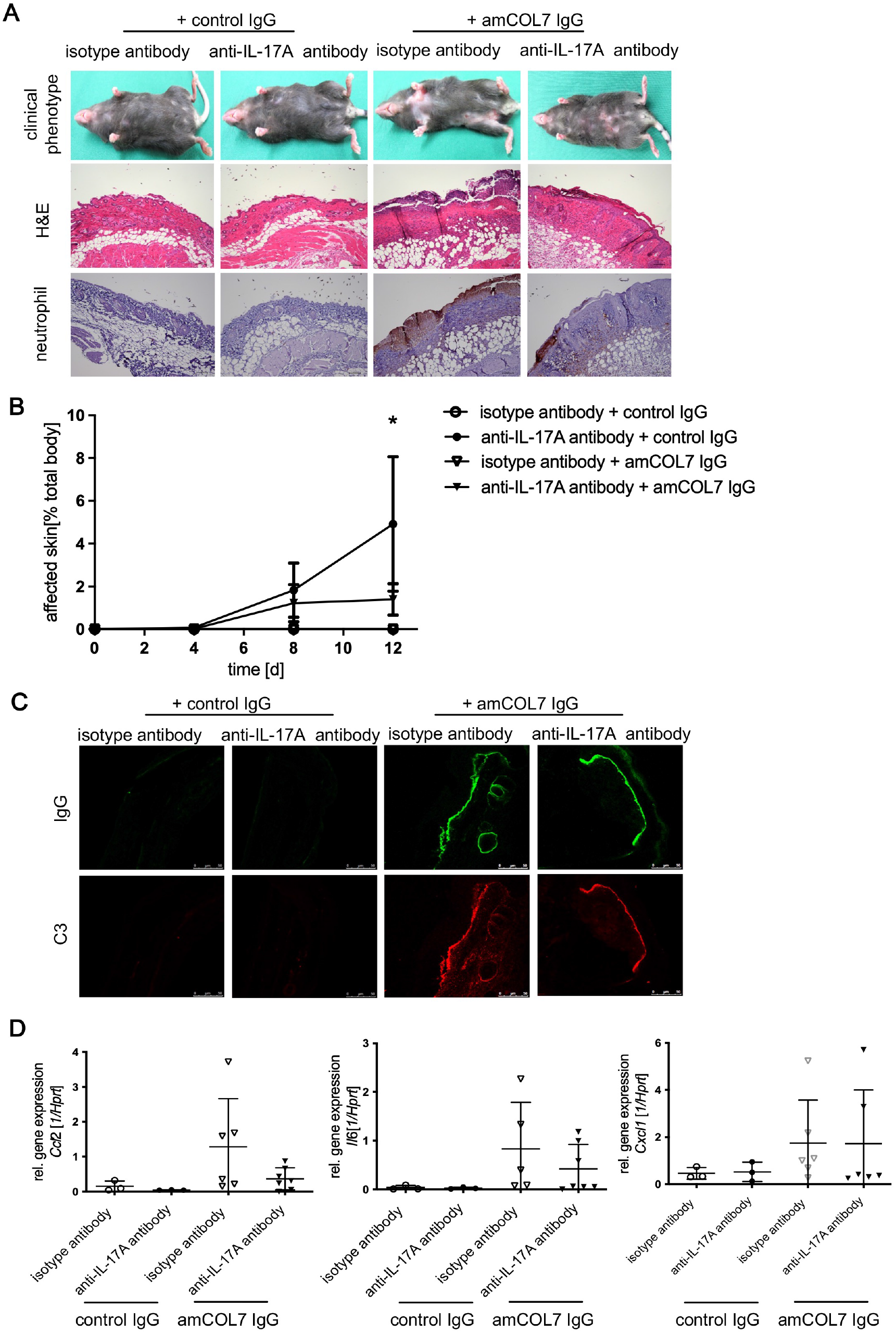
Pharmacological inhibition of IL-17A attenuates antibody transfer-induced EBA. C57BL/6 mice received six subcutaneous injections of anti-mCOL7 IgG (n = 5) or corresponding amounts of normal rabbit IgG (n = 5) every other day. To neutralize IL-17A, mice were treated with an anti-IL-17A antibody or the corresponding isotype control antibody on days −2, 0, 2, 4, 6, 8, and 10. A: Representative clinical photographs and histological analysis of lesional skin. Formalin-fixed ear skin sections were stained with hematoxylin and eosin (H&E), and neutrophil infiltration was assessed by immunohistochemistry using an anti-neutrophil antibody. Scale bars = 100 µm. B: Clinical disease severity expressed as percentage of affected body surface area over time. C: Tissue-bound IgG and complement C3 deposition at the dermal–epidermal junction visualized by direct immunofluorescence microscopy. D: Quantification of mRNA expression of selected inflammatory cytokines and chemokines in lesional skin by quantitative real-time PCR. Data are shown as mean ± SD (n = 3-5 per group). *p < 0.05, **p < 0.01; Mann–Whitney rank-sum test.

### γδ T cells are the main source of IL-17A in experimental EBA

To identify the cellular source of IL-17A in EBA, IL-17A-eGFP mice were subjected to antibody transfer-induced EBA by injection of anti-mCOL7 IgG. Following anti-mCOL7 IgG transfer, IL-17A-eGFP mice developed EBA comparable to that observed in C57BL/6 mice, with erosions, erythema, and hair loss on the ears and body surface (Figure 2A). Histological analysis revealed marked cellular infiltration in both C57BL/6 and IL-17A-eGFP mice after anti-mCOL7 IgG transfer (Figure 2A). Quantification of overall disease severity based on the affected body surface area showed no difference between IL-17A-eGFP and C57BL/6 mice (Figure 2B). To define the IL-17A-producing cells in experimental EBA, cells were isolated from lesional skin of IL-17A-eGFP mice, stained for cell surface markers, and analyzed by flow cytometry. Among IL-17A^+^ cells, γδ T cells were identified as the major cellular source of IL-17A (62.98 ± 12.33%), followed by CD4^+^ T cells (29.95 ± 11.42%) and Ly6G^+^ neutrophils (9.36 ± 6.88%) (Figure 2C, D). Collectively, these data establish γδ T cells as the dominant source of IL-17A in experimental EBA and implicate this population as a critical cellular component of the pathogenic inflammatory response.

**Figure 2.**
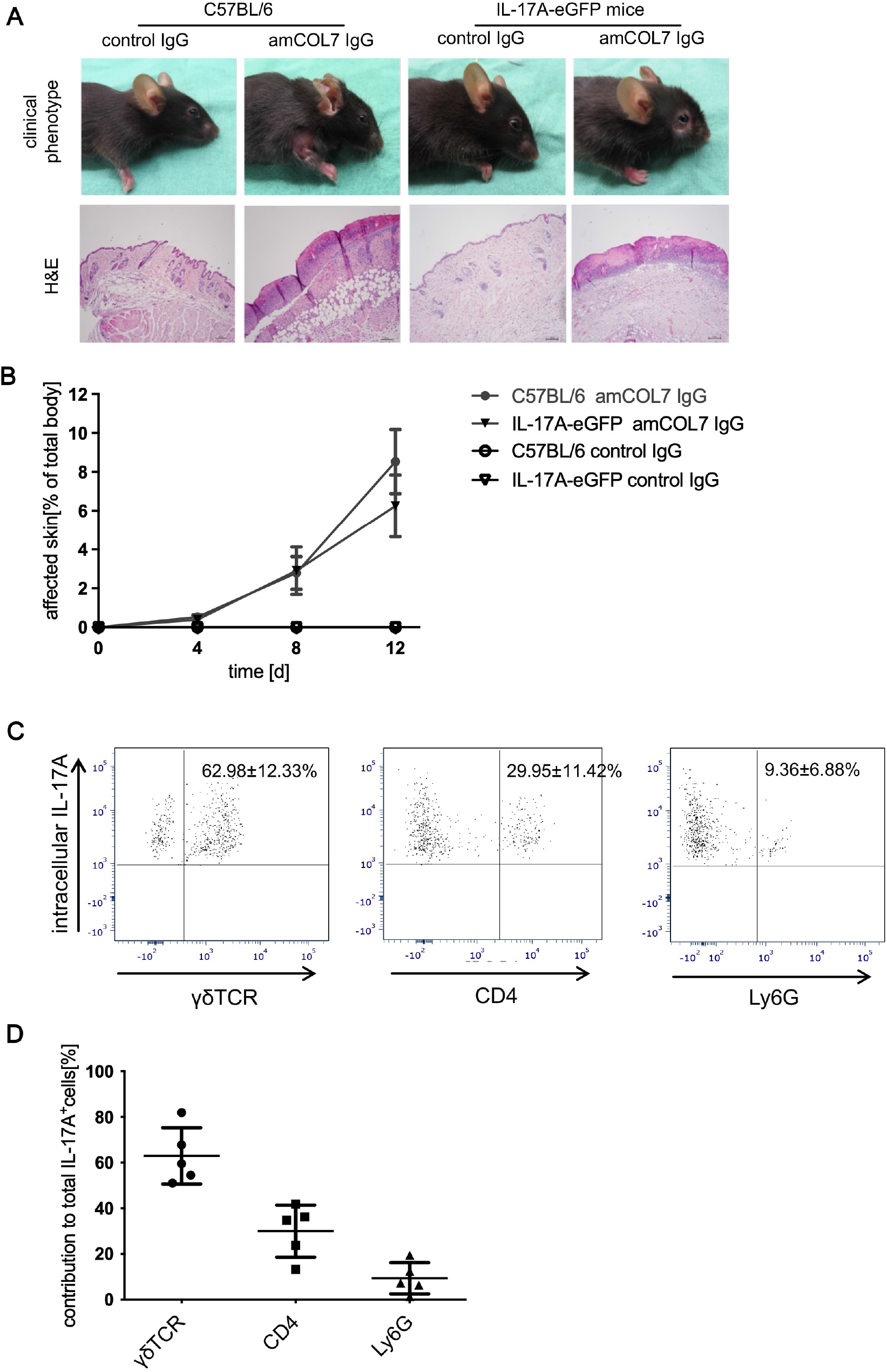
γδ T cells are the main source of IL-17A in antibody transfer-induced EBA. IL-17A-eGFP reporter mice received six subcutaneous injections of anti-mCOL7 IgG or corresponding amounts of normal rabbit IgG every other day. A: Representative clinical photographs and H&E-stained ear skin sections from C57BL/6 and IL-17A-eGFP reporter mice. Scale bar = 100 µm. B: Clinical disease severity expressed as percentage of affected body surface area over time. C, D: Flow cytometric analysis of IL-17A-producing cells isolated from lesional ear skin of anti-mCOL7 IgG-treated IL-17A-eGFP reporter mice. Cells were stained for CD45, CD4, γδ TCR, Ly6G, and viability markers and analyzed by flow cytometry. C: Representative gating of live single IL-17A+ cells according to γδ TCR, CD4, and Ly6G expression. D: Quantification of the relative contribution of γδ T cells, CD4+ T cells, and Ly6G+ neutrophils among live single IL-17A+ cells. Data are shown as mean ± SD (n = 5 per group). *p < 0.05, **p < 0.01; Mann–Whitney rank-sum test.

### γδTCR^-/-^ mice are protected from the pathogenic effects of anti-mCOL7 IgG

To determine the contribution of IL-17A-producing γδ T cells to EBA, experimental EBA was induced in γδTCR^-/-^ mice, and IL-17A levels and disease severity were evaluated. After anti-mCOL7 IgG injection, numerous IL-17A^+^ cells were detected in the skin of C57BL/6 mice, whereas the frequency of IL-17A^+^ cells was markedly reduced in γδTCR^-/-^ mice (2.272 ± 1.332% *vs* 0.968 ± 0.410%, p < 0.05, Figure 3A). Consistently, mRNA levels of IL-17A and IL-17F in affected skin were also significantly reduced in γδTCR^-/-^ mice compared with C57BL/6 mice (Figure 3B,C).

**Figure 3.**
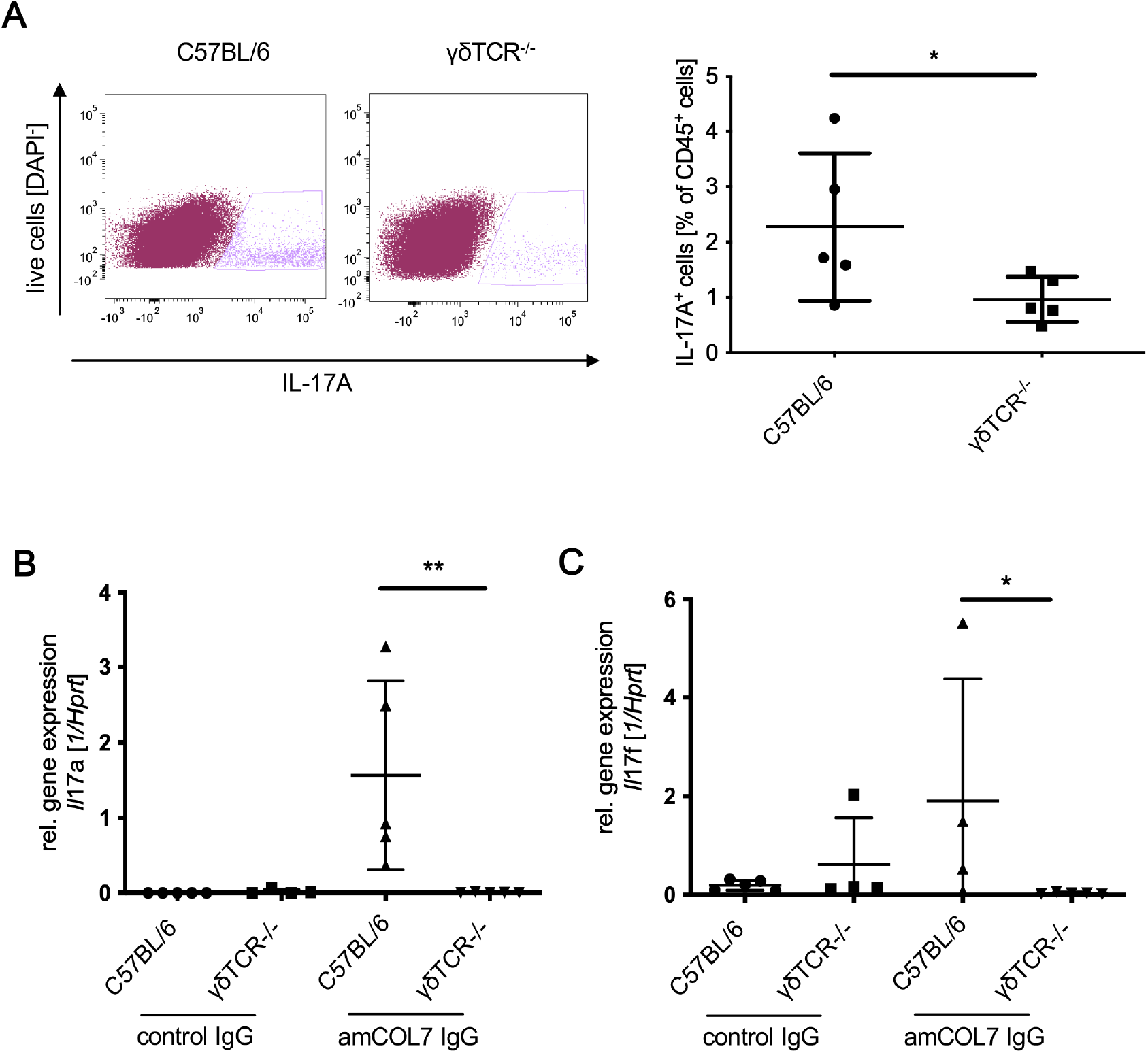
γδ TCR deficiency reduces IL-17A levels in lesional skin during antibody transfer-induced EBA. Cells were isolated from lesional skin of C57BL/6 and γδ TCR^−/−^ mice 12 days after anti-mCOL7 IgG injection and stimulated with PMA/ionomycin in the presence of GolgiPlug. Cells were stained for surface markers (CD45, CD3, CD4, Ly6G, γδ TCR, and B220), followed by intracellular staining for IL-17A. A: Representative flow cytometric analysis of IL-17A+ leukocytes and quantification of IL-17A-producing cells in lesional skin of C57BL/6 and γδ TCR^−/−^ mice. B, C: mRNA expression of Il17a (B) and Il17f (C) in skin from C57BL/6 and γδ TCR^−/−^ mice on day 12 after injection of normal rabbit IgG or anti-mCOL7 IgG, determined by quantitative real-time PCR. Data are shown as mean ± SD (n = 4-5 per group). *p < 0.05, **p < 0.01; Mann–Whitney rank-sum test.

Furthermore, after anti-mCOL7 IgG injection, C57BL/6 mice developed severe skin blistering, whereas γδTCR^-/-^ mice showed a milder EBA phenotype (Figure 4A). Comparison of clinical scores between both strains on days 4, 8, and 12 revealed significantly reduced affected body surface area in γδTCR^-/-^ mice at all time points (day 4: 0.472 ± 0.135% vs 0.245 ± 0.084%, p < 0.05; day 8: 3.756 ± 1.621% vs 1.305 ± 0.322%, p < 0.05; day 12: 7.325 ± 3.165% vs 3.305 ± 1.138%, p < 0.01, in WT and γδTCR^-/-^ mice, respectively; Figure 4B). The clinical improvement was accompanied by reduced dermal inflammatory infiltrates in γδTCR^-/-^ mice compared with control mice. Accordingly, mRNA expression of the pro-inflammatory cytokine IL-1β was decreased in γδTCR^-/-^ mice compared with C57BL/6 mice after anti-mCOL7 IgG injection (Figure 4C). mRNA levels of the neutrophil-associated chemokines *Cxcl2* and *Ccl2* were also reduced, although the differences did not reach statistical significance compared with C57BL/6 mice (Figure 4C). In contrast, levels of tissue-bound IgG and C3 were similar in all mice (Figure 4D), indicating that the reduced clinical severity was not attributable to altered IgG metabolism or impaired complement deposition. Together, these results indicate that γδ T cells promote EBA pathogenesis by driving IL-17A-dependent inflammatory responses downstream of autoantibody binding and complement deposition.

**Figure 4.**
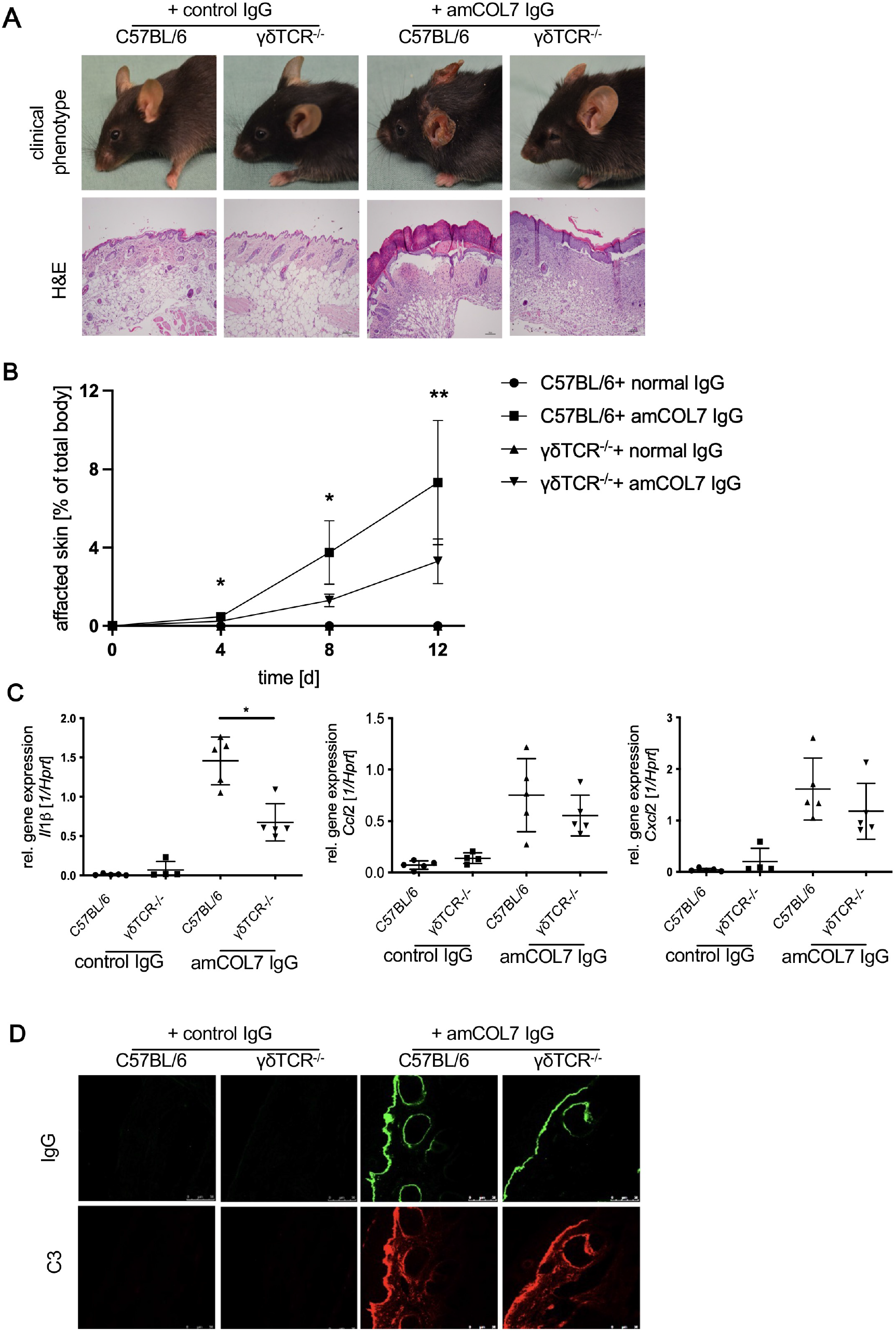
γδ TCR^−/−^ mice are protected from antibody transfer-induced EBA. C57BL/6 and γδ TCR^−/−^ mice received six subcutaneous injections of anti-mCOL7 IgG (n = 5) or corresponding amounts of normal rabbit IgG (n = 5) every other day. A: Representative clinical photographs and H&E-stained ear skin sections from C57BL/6 and γδ TCR^−/−^ mice. Scale bar = 100 µm. B: Clinical disease severity expressed as percentage of affected body surface area over time. C: Quantification of mRNA expression of selected inflammatory cytokines and chemokines in lesional skin by quantitative real-time PCR. D: Tissue-bound IgG and complement C3 deposition at the dermal–epidermal junction visualized by direct immunofluorescence microscopy. Data are shown as mean ± SD (n = 4-5 per group). *p < 0.05, **p < 0.01; Mann–Whitney rank-sum test.

### IL-1β promotes IL-17A production by γδ T cells *in vitro*

Given the dominant contribution of γδ T cells to IL-17A production in experimental EBA, we next sought to identify the upstream signals that trigger IL-17A secretion by this cell population. To this end, an *in vitro* assay was performed to identify mechanisms inducing IL-17A secretion by γδ T cells. After isolation from γδTCR-eGFP mice, γδ T cells were stimulated with mCOL7 immune complexes, C3a, IL-23, and IL-1β, as indicated, for 72 h (Figure 5). γδ T cells did not produce IL-17A in response to anti-COL7 immune complexes or C3a alone. In contrast, co-stimulation with IL-1β significantly increased IL-17A production by γδ T cells. Stimulation with IL-23 together with immune complexes induced only a slight increase in IL-17A levels in γδ T-cell supernatants. Notably, even in the absence of immune-complex stimulation, IL-1β strongly promoted IL-17A production by γδ T cells, whereas IL-23 did not. ollectively, these findings suggest that IL-1β is a potent inducer of IL-17A production by γδ T cells and may serve as an important upstream regulator of pathogenic IL-17A responses in experimental EBA.

**Figure 5.**
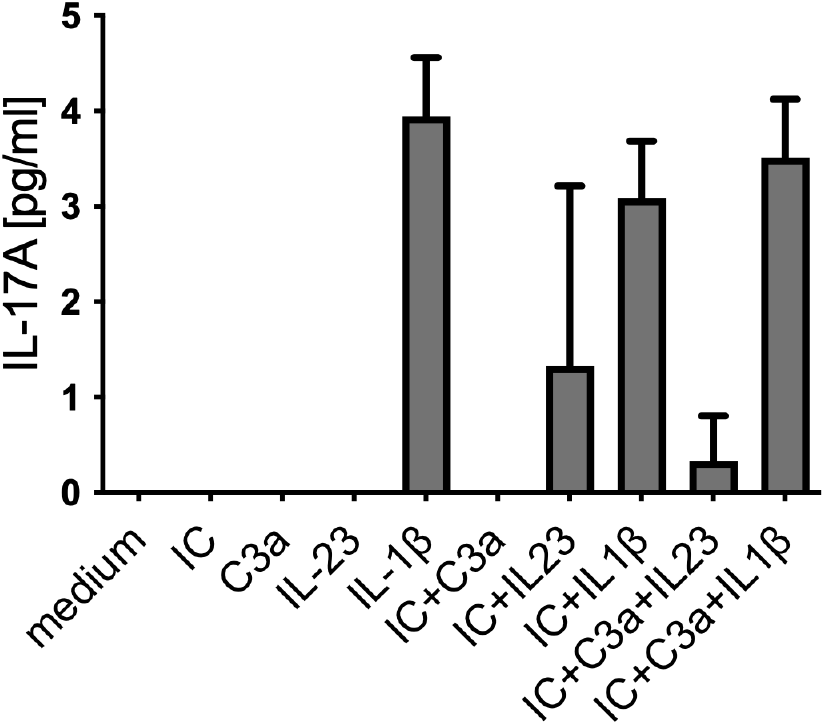
IL-1β promotes IL-17A production by γδ T cells in vitro. γδ T cells were isolated from spleens of γδ TCR-eGFP mice by fluorescence-activated cell sorting after exclusion of non-γδ T-cell populations using antibodies against CD4, CD8, CD11b, B220, and NK1.1. Sorted γδ T cells were stimulated with immobilized mCOL7 immune complexes, C3a, IL-23, and IL-1β, as indicated, for 72 h. IL-17A concentrations in culture supernatants were quantified by ELISA. Data are shown as mean ± SD from three independent experiments, each performed in duplicate.

### EBA severity is not alleviated in IL-23p19^-/-^ mice

Since IL-23 is considered a central regulator of pathogenic IL-17 responses in several inflammatory and autoimmune diseases (Schinocca et al., 2021), we next assessed whether the classical IL-23/IL-17 axis contributes to EBA pathogenesis *in vivo*, despite the strong IL-1β-dependent induction of IL-17A observed in our *in vitro* experiments. To this end, functional relevance of the IL-23/IL-17A axis in experimental EBA was analyzed in IL-23p19^-/-^ mice. After repeated administration of pathogenic anti-mCOL7 IgG, disease severity was assessed on days 4, 8, and 12. IL-23p19^-/-^ mice developed clinical disease comparable to that of control mice throughout the experiment (day 12: 6.518 ± 1.690% vs 3.927 ± 1.631%, p > 0.05, Figure 6A). Consistently, clinical phenotype, histopathological changes, and neutrophil infiltration in skin lesions did not differ between control and IL-23p19^-/-^ mice after antibody transfer (Figure 6B). mRNA levels of IL-17A and IL-17F in affected skin were reduced in IL-23p19^-/-^ mice compared with C57BL/6 mice but did not differ significantly. Similarly, mRNA expression of associated inflammatory cytokines and chemokines, including *Il6*, *Cxcl1*, and *Ccl2*, was comparable between IL-23p19^-/-^ and C57BL/6 mice (Figure 6C). Taken together, these findings suggest that the pathogenic IL-17A response in experimental EBA is not critically dependent on IL-23, prompting us to further investigate the contribution of IL-1 signaling *in vivo*.

**Figure 6.**
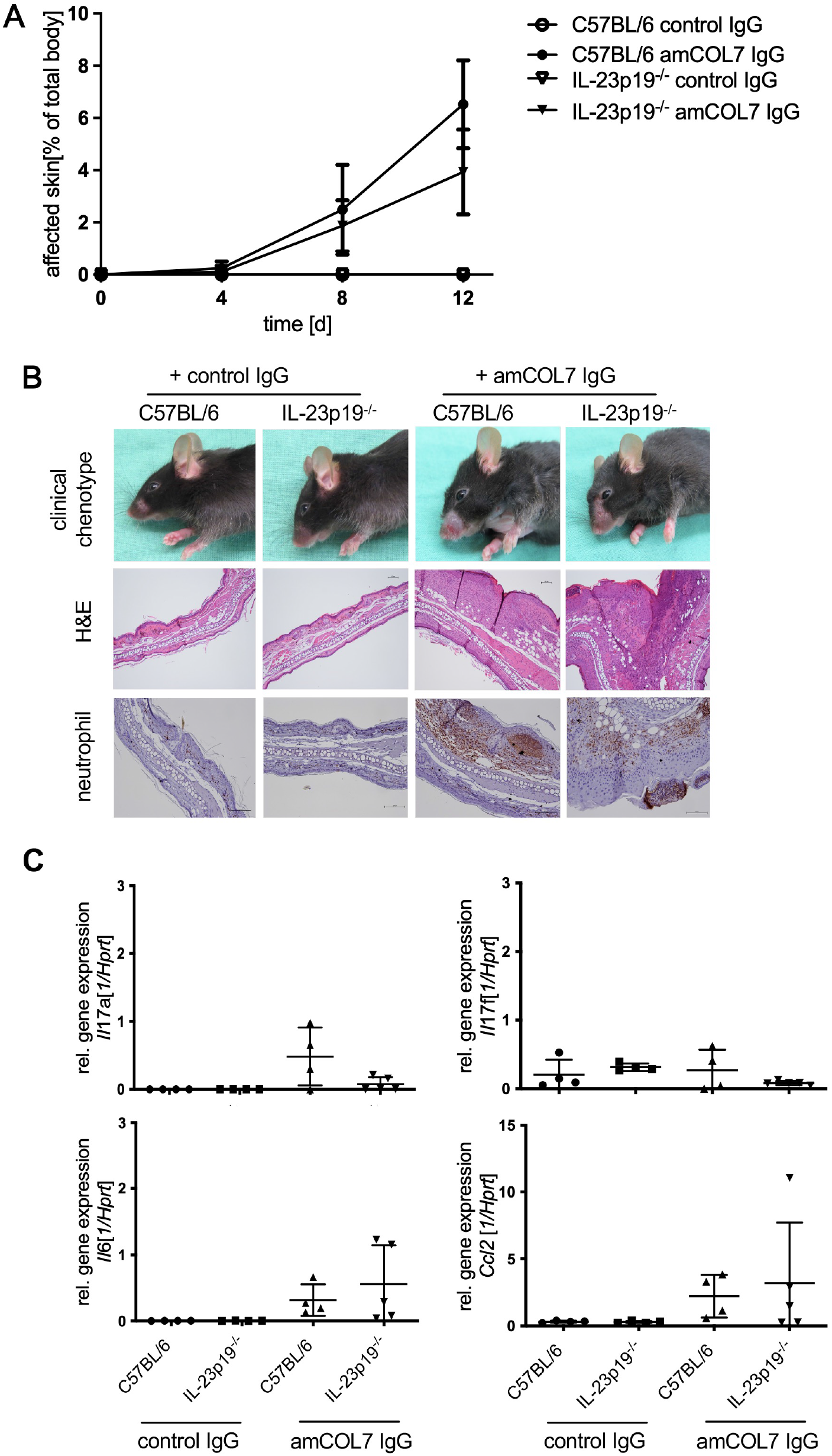
IL-23p19 deficiency does not attenuate antibody transfer-induced EBA. C57BL/6 and IL-23p19^−/−^ mice received six subcutaneous injections of anti-mCOL7 IgG (n = 5) or corresponding amounts of normal rabbit IgG (n = 5) every other day. A: Clinical disease severity expressed as percentage of affected body surface area over time. B: Representative clinical photographs, H&E-stained ear skin sections, and immunohistochemical detection of neutrophil infiltration in lesional skin. Scale bars = 100 µm. C: Quantification of mRNA expression of selected inflammatory cytokines and chemokines in lesional skin by quantitative real-time RT-PCR. Data are shown as mean ± SD (n = 4–5 per group). **p < 0.01, ***p < 0.001; Mann–Whitney rank-sum test.

### Antibody-induced EBA is attenuated in IL-1R^-/-^ mice

To determine whether IL-1 signaling is required for IL-17A production during EBA development, experimental EBA was induced in IL-1R^-/-^ and C57BL/6 mice. Mice of both strains developed experimental EBA; however, the overall extent of skin blistering was markedly reduced in IL-1R^-/-^ mice compared with control mice throughout the observation period (day 4: 2.078 ± 1.171% vs 0.867 ± 0.419%, p < 0.01; day 8: 7.978 ± 2.861% vs 3.578 ± 1.484%, p < 0.005; day 12: 16.578 ± 5.671% vs 8.178 ± 2.563%, p < 0.001, in C57BL/6 and IL-1R^-/-^ mice, respectively; Figure 7A). The clinical phenotype and histopathological alterations are shown in Figure 7B. Following transfer of anti-mCOL7 IgG, both C57BL/6 and IL-1R-/-mice developed clinical signs of EBA; however, disease manifestations were markedly less severe in IL-1R^-/-^ mice. Compared with C57BL/6 mice, IL-1R^-/-^ mice exhibited reduced hair loss, erythema, crust formation and skin blistering. Histological examination of lesional skin confirmed the attenuated disease phenotype, revealing reduced epidermal disruption and less pronounced inflammatory cell infiltration in IL-1R^-/-^ mice. To further assess the inflammatory response, expression levels of selected cytokines and chemokines were quantified in affected skin (Figure 7C). Consistent with the reduced clinical and histopathological severity, IL-1R^-/-^ mice showed lower expression of several inflammatory mediators, with *Ccl2* being significantly decreased compared to C57BL/6 control mice. In addition, *ll17a* mRNA levels were reduced in lesional skin of IL-1R^-/-^ mice although this difference did not reach statistical significance. Together, these findings indicate that IL-1 signaling contributes to the amplification of local inflammatory responses and promotes tissue damage during antibody transfer-induced EBA.

**Figure 7.**
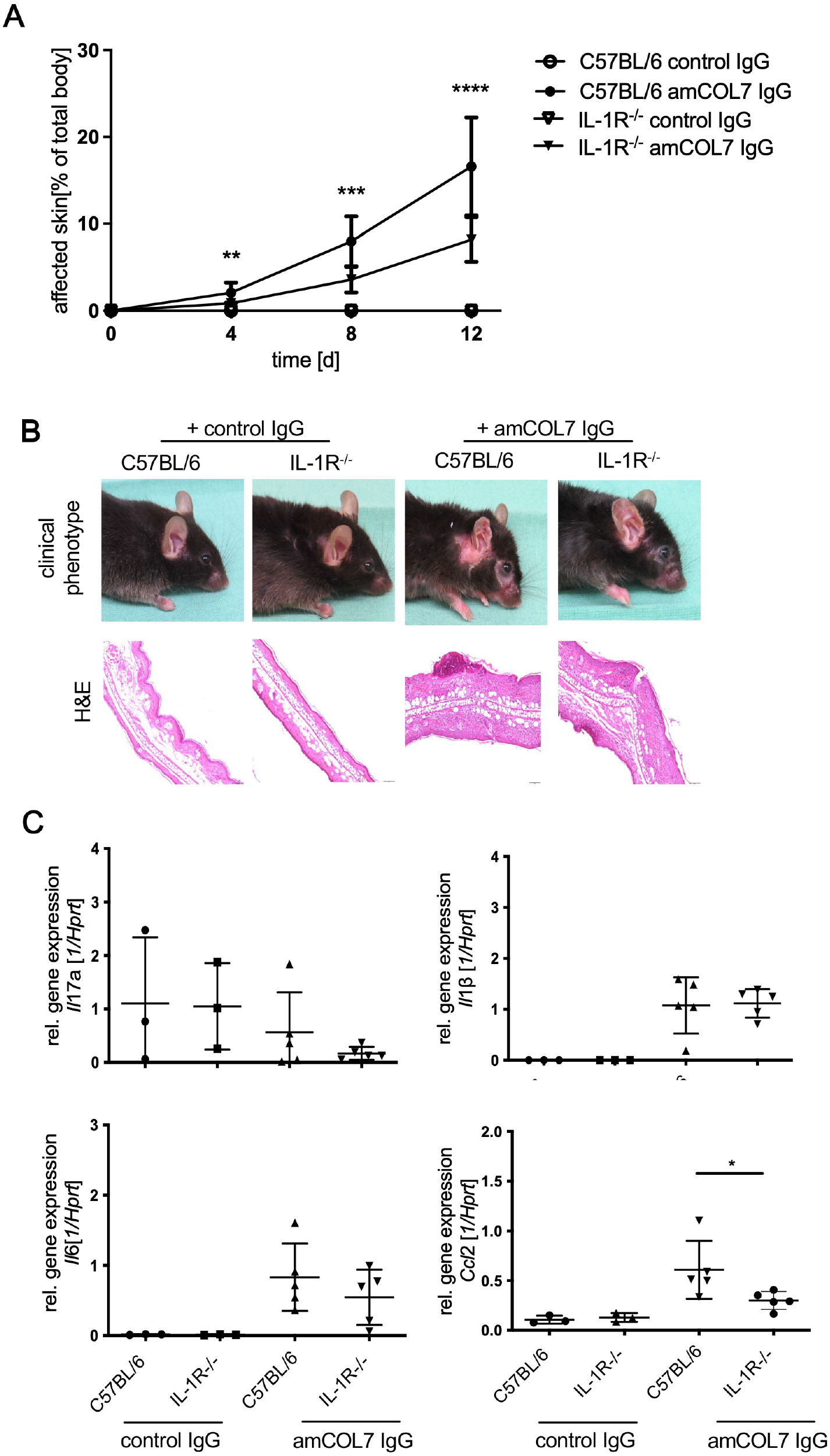
IL-1 signaling is required for full development of antibody transfer-induced EBA. C57BL/6 and IL-1R^−/−^ mice received six subcutaneous injections of anti-mCOL7 IgG (n = 9) or corresponding amounts of normal rabbit IgG (n = 5) every other day. A: Clinical disease severity expressed as percentage of affected body surface area over time. B: Representative clinical photographs and H&E-stained ear skin sections from C57BL/6 and IL-1R^−/−^ mice. Scale bar = 100 µm. C: Quantification of mRNA expression of selected inflammatory cytokines and chemokines in lesional skin by quantitative real-time RT-PCR. Data are shown as mean ± SD (n = 3–5 per group). *p < 0.05, **p < 0.01, ***p < 0.001; Mann– Whitney rank-sum test.

## DISCUSSION

Although a pathogenic role of IL-17A has been implicated in several autoimmune diseases, whether and how IL-17A contributes to the development of EBA has remained unclear. In the present study, we identify IL-17A as a pathogenic mediator in EBA and demonstrate that γδ T cells, rather than conventional Th17 cells, constitute the predominant cellular source of IL-17A during disease development. Furthermore, our findings indicate that IL-1β, but not IL-23, is the principal upstream cytokine driving IL-17A production by γδ T cells, consistent with our *in vivo* data. Specifically, IL-23p19^-/-^ mice showed no reduction in EBA severity, whereas IL-1R deficiency markedly ameliorated disease. Together, these findings define a previously unrecognized IL-1β/γδ T cell/IL-17A axis that contributes to EBA pathogenesis.

IL-17A functions as a major regulator of neutrophil-dominated inflammation by inducing chemokine production, promoting granulopoiesis, and activating tissue-resident stromal and epithelial cells (Liang et al., 2006; Ye et al., 2001). The pathogenic role of IL-17A in EBA was supported by pharmacological evidence. Therapeutic neutralization of IL-17A significantly reduced disease progression in wild-type animals. This result strongly supports a direct pathogenic role for IL-17A. These findings are consistent with studies in other autoimmune diseases, including psoriasis, psoriatic arthritis, and ankylosing spondylitis, in which IL-17A has emerged as a central inflammatory mediator (Cai et al., 2011; Zhu and Qian, 2012) and an effective therapeutic target (Baeten et al., 2013; McInnes et al., 2025; McInnes et al., 2015). Importantly, our results also align with evidence demonstrating that IL-17A is functionally relevant and represents a potential therapeutic target in BP, another major subepidermal blistering disease (Chakievska et al., 2019). Thus, our findings extend the spectrum of IL-17A-driven autoimmune blistering diseases and suggest that IL-17A may represent a promising therapeutic target in EBA.

A notable finding of this study is that γδ T cells were identified as the dominant source of IL-17A in experimental EBA. Although Th17 cells are generally considered major producers of IL-17A in many autoimmune diseases (Chakievska et al., 2019; Komiyama et al., 2006; van der Fits et al., 2009), increasing evidence indicates that tissue-resident γδ T cells are important contributors to IL-17A-mediated inflammation in barrier tissues, particularly in the skin (Cai et al., 2011; Chojnacka-Purpurowicz et al., 2024; Martin et al., 2009). Unlike conventional adaptive T cells, γδ T cells possess innate-like properties that enable rapid cytokine production in response to tissue stress signals without requiring extensive antigen-specific clonal expansion (Bonneville et al., 2010; Khairallah et al., 2018). This functional feature positions γδ T cells as important regulators of early inflammatory responses after tissue injury. Consistent with this concept, γδ T cells accumulated in EBA lesions, and a substantial proportion of infiltrating γδ T cells expressed IL-17A. Moreover, genetic deficiency of γδ T cells significantly reduced disease severity, inflammatory cell recruitment, and local IL-17A expression, demonstrating that γδ T cells are required for the full development of experimental EBA. These results indicate that γδ T cells are not merely bystander inflammatory cells but represent a critical cellular component initiating pathogenic IL-17A responses in antibody-mediated skin injury. The preferential involvement of γδ T cells in EBA highlights the importance of tissue-specific immune regulation in autoimmune blistering diseases.

Skin-resident γδ T cells constitute a specialized immunosurveillance network that continuously monitors epithelial integrity and rapidly responds to tissue damage (Ribot et al., 2021). Among these populations, dermal Vγ4^+^ and Vγ6^+^ γδ T cells have been identified as major sources of IL-17A in inflammatory skin conditions, whereas epidermal dendritic epidermal T cells primarily contribute to tissue homeostasis and repair (Cai et al., 2011; Fleißner et al., 2025; Nielsen et al., 2017). Although the γδ T-cell subsets responsible for IL-17A production were not further characterized in this study, our findings suggest that IL-17-producing dermal γδ T cells may represent the dominant pathogenic population during antibody-induced EBA. Further characterization of the γδ T-cell subsets involved in EBA may provide additional mechanistic insight.

Interestingly, the cellular source of IL-17A appears to differ among autoimmune blistering diseases. In BP, several immune-cell populations, including neutrophils, macrophages, and activated T cells, have been proposed as potential sources of IL-17A (Chakievska et al., 2019; Le Jan et al., 2014). In contrast, our data identify γδ T cells as the predominant IL-17A-producing population in experimental EBA. This difference may reflect disease-specific inflammatory milieus shaped by distinct autoantigens, tissue compartments, and modes of immune activation. Thus, autoantibody specificity alone may not fully determine the resulting disease phenotype; rather, local cytokine networks and the composition of tissue-resident immune cells are likely to critically shape inflammatory outcomes.

By identifying γδ T cells as the dominant source of pathogenic IL-17A, our study provides a new cellular framework for understanding how autoantibody deposition initiates and amplifies inflammatory tissue damage in EBA. Another important observation is that IL-17A production in experimental EBA is primarily controlled by IL-1β rather than by the classical IL-23-dependent pathway. The IL-23/IL-17 axis is widely recognized as a central driver of chronic autoimmune inflammation (Schinocca et al., 2021; Xue et al., 2014). In diseases such as experimental autoimmune encephalomyelitis, inflammatory bowel disease, and psoriasis, IL-23 promotes the expansion, stabilization, and pathogenic differentiation of IL-17-producing T cells, thereby sustaining inflammatory responses (Gaffen et al., 2014). However, our data demonstrate that IL-23 deficiency does not significantly affect disease severity in experimental EBA, indicating that IL-23 is not required for disease-associated IL-17A induction in this antibody-mediated model. Instead, IL-17A appears to be induced through an alternative regulatory mechanism. This lack of requirement for IL-23 dependence is particularly notable because IL-23 is considered essential for maintaining pathogenic Th17 responses in many autoimmune diseases. Unlike chronic antigen-driven autoimmune disorders, the antibody-transfer model of EBA represents a rapidly initiated effector-phase response triggered by deposition of pathogenic anti-COL7 antibodies at the dermal–epidermal junction. This process induces immediate complement activation, Fc receptor signaling, and inflammatory mediator release (Ludwig and Zillikens, 2011). Under these conditions, tissue-resident γδ T cells may respond directly to inflammatory cytokines generated within the local microenvironment, allowing rapid IL-17A production without requiring the need for IL-23-dependent differentiation or expansion.

Our findings are consistent with previous studies showing that IL-1 signaling plays a crucial role in regulating IL-17-producing lymphocytes. IL-1β promotes the differentiation of pathogenic IL-17-producing T cells and is required for autoimmune inflammation in models such as experimental autoimmune encephalomyelitis (Chung et al., 2009; Lalor et al., 2011; Sutton et al., 2006). In line, IL-1β-deficient mice have previously been shown to develop significantly less skin lesions upon injection of IgG against COL7 and COL17, the major autoantigen of BP, respectively. Furthermore, elevated levels of IL-1β have been found the blister fluid and skin of BP patients (Heidari et al., 2025; Holtsche et al., 2020; Sadeghi et al., 2015). Moreover, IL-1β can directly stimulate γδ T cells to produce IL-17A, often acting together with IL-23 to amplify innate immune responses (Martin et al., 2009; Sutton et al., 2009). In the present study, IL-1β directly induced IL-17A production by γδ T cells *in vitro*, whereas IL-23 alone had only limited effects. Furthermore, IL-1R^-/-^ mice were markedly protected from EBA development, supporting the *in vivo* relevance of this pathway.

Mechanistically, IL-1β is known to promote IL-17A expression through induction of the transcription factor RORγt, which is essential for IL-17-producing lymphocytes (Chung et al., 2009). Similar IL-1β-dependent activation of γδ T cells has been reported in cancer immunology and experimental autoimmune encephalomyelitis (Sutton et al., 2006; Sutton et al., 2009). Although classical models suggest that IL-1β and IL-23 often cooperate to promote innate IL-17 production from γδ T cells (Sutton et al., 2009), our findings reveal a distinct, IL-23-independent dominance of IL-1β in this autoantibody-induced inflammatory setting.

We propose that the dispensability of IL-23 reflects the rapid initiation kinetics of the EBA antibody-transfer model. Passive transfer of anti-type VII collagen antibodies triggers an immediate, complement-dependent inflammatory cascade. Under these conditions, IL-1β may be rapidly processed and released by activated resident or recruited cells, providing an early and sufficient stimulus for pre-committed, tissue-resident γδ T cells. Notably, this early cytokine response may also involve alternative, caspase-1-independent pathways of IL- 1β release that directly promote blister formation during autoantibody-induced tissue injury (Sadeghi et al., 2015; Sutton et al., 2006). Rapidly available IL-1β may then activate skin-resident γδ T cells to produce pathogenic IL-17A, whereas IL-23 may function primarily as an amplifying rather than initiating signal. Collectively, these observations support a pathogenic pathway in EBA in which IL-1β-driven activation of γδ T cells leads to IL-17A production, neutrophil recruitment, and subsequent tissue damage.

Several limitations should be acknowledged. First, all experiments were performed in passive antibody-transfer models, which primarily reflect the effector phase of EBA and do not address the potential role of IL-17A in autoantibody generation or loss of immune tolerance. Therefore, the relevance of γδ T-cell-derived IL-17A in human EBA remains to be established. Future studies using immunization-induced models of EBA may help determine whether this pathway also contributes to disease initiation. In addition, the specific γδ T-cell subset responsible for IL-17A production, the cellular source of IL-1β within EBA lesions, and the target cells responding to IL-17A should be defined in future studies.

Despite these limitations, our findings have important translational implications. We identify IL-17A as a critical pathogenic mediator in experimental EBA and demonstrate that disease development is largely driven by an IL-1β-dependent γδ T-cell response. Therapeutic targeting of IL-17A has shown substantial clinical efficacy in psoriasis and other chronic inflammatory diseases (Brembilla et al., 2018), whereas IL-1β-directed therapies, including anakinra and canakinumab, have proven beneficial in several autoinflammatory diseases (Broderick and Hoffman, 2022). The identification of the IL-1β/γδ T cell/IL-17A axis as acentral pathogenic pathway in EBA suggest that blockade of either IL-17A or IL-1β may represent a promising therapeutic approach for this disease. Future studies should determine whether IL-17A-producing γδ T cells are similarly enriched in skin lesions and peripheral blood of patients with EBA and assess the therapeutic potential of IL-17A- and IL-1β-targeted biologics in translational and clinical settings.

In conclusion, this study identifies IL-17A as a pivotal driver of experimental EBA and establishes γδ T cells as its predominant cellular source. In contrast to many autoimmune disorders that depend on the classical IL-23/TH17 pathway, EBA appears to be characterized by an IL-1β-driven γδ T-cell response that promotes pathogenic IL-17A production. These findings provide new insight into the immunopathogenesis of EBA immunopathogenesis and highlight the IL-1β/γδ T cell/IL-17A axis as a potential therapeutic target for this severe autoimmune blistering disease.

## MATERIALS AND METHODS

### Mice

*Il23a*^tm1Ngh/tm1Ngh^ (IL-23p19^-/-^) (Ghilardi et al., 2004) mice were bred at the Research Center Borstel (Borstel, Germany). *Tcrd*^tm1Mom^ (γδTCR^-/-^) (Itohara et al., 1993) and *Trdc*^tm1Mal^ (γδTCR-eGFP) (Prinz et al., 2006) mice were obtained from the Institute of Immunology at Hannover Medical School (Hannover, Germany), *Il1r1*^tm1Imx^ (IL-1R^-/-^) (Glaccum et al., 1997) mice were provided by the Institute of Experimental Dermatology at the University of Lübeck (Lübeck, Germany) and C57BL/6J mice were purchased from Charles River Laboratories (Sulzfeld, Germany). All animals used in the experiments were females and 8–16 weeks old. Mice were housed under specific pathogen-free conditions with a 12-hour light/dark cycle at the animal facility of the Research Center Borstel. All animal experiments were reviewed and approved by the Animal Research Ethics Board of the Ministry of Agriculture, Rural Areas, European Affairs and Consumer Protection of the State of Schleswig-Holstein, Kiel, Germany (approval numbers 91-9/10, 4-1/13, 12-2/14, and 109-10/19). Clinical examinations, biopsies, and blood sampling were performed under anesthesia using intraperitoneal administration of ketamine (100 μg/g) and xylazine (15 μg/g).

### Anti-mCOL7 antibodies

Rabbit anti-mCOL7c antibodies and rabbit control IgG were prepared as described previously (Sitaru et al., 2005). Briefly, New Zealand White rabbits were immunized subcutaneously with recombinant mCOL7c protein (amino acid residues 757–967) emulsified in Freund’s complete adjuvant. Animals were boosted twice with mCOL7c protein emulsified in Freund’s incomplete adjuvant. Sera from immunized rabbits were collected and used for isolation of total IgG by Protein G Sepharose Fast Flow affinity column chromatography. Control rabbit IgG was isolated from sera of non-immunized rabbits. The reactivity of all IgG fractions was analyzed by immunofluorescence microscopy using murine skin.

### Induction and evaluation of experimental EBA

Experimental EBA was induced as previously described (Sitaru et al., 2005), with minor modifications. Female mice aged 8–16 weeks received six subcutaneous injections of 2–3 mg rabbit anti-mCOL7 IgG, with exact doses specified in the respective experimental protocols, or normal rabbit IgG into the abdomen every other day on days 0, 2, 4, 6, 8, and 10. Disease severity was expressed as the percentage of affected body surface area and assessed on days 4, 8, and 12. For IL-17A blockade, mice received anti-IL-17A antibody (clone 17F3, BioXCell; 200 μg/mouse) every other day from day −2 to day 10, for a total of seven injections. Control mice received rat IgG1 isotype antibody (clone MOPC-21, BioXCell; 200 μg/mouse, i.p.) according to the same schedule. Cutaneous lesions, crusts, erythema, and alopecia were scored for each body region as follows: 2.5% each for the ears, snout, and oral mucosa; 0.5% for each eye; 9% for the head and neck, excluding eyes, ears, snout, and oral mucosa; 5% for each forelimb; 10% for the hind limbs and tail; and 40% for the trunk (Gross et al., 2024). The sum of these values yielded the overall clinical score. On day 12, mice were sacrificed, and sera and biopsies were collected.

### Hematoxylin / eosin staining of lesional skin

For histological analysis, skin and ear samples were fixed in 4% buffered formaldehyde and embedded in paraffin. Paraffin sections of 5 μm thickness were prepared and stained with hematoxylin and eosin (H&E) according to standard protocols for histological evaluation.

### Immunohistochemical detection of neutrophil infiltration

Formalin-fixed, paraffin-embedded skin sections were deparaffinized in xylene and rehydrated through graded ethanol. Antigen retrieval was performed by heating sections at 121 °C in 10 mM citrate buffer (pH 6.0) for 1 h. Endogenous peroxidase activity, endogenous biotin, and nonspecific binding were blocked using 3% H_2_O_2_, biotin blocking solution (Vector Laboratories), and 5% BSA, respectively. Sections were incubated overnight at 4 °C with anti-murine neutrophil antibody (07-APR, Cedarlane), followed by incubation with biotinylated secondary antibody (Jackson ImmunoResearch) for 45 min at room temperature, avidin-biotinylated HRP (Vector Laboratories) for 20 min, and DAB (Vector Laboratories) for visualization. Sections were counterstained with hematoxylin for 5 min, and images were acquired by bright-field microscopy (Nikon).

### Detection of tissue-bound IgG and complement

Tissue-bound IgG and complement C3 deposition were detected by immunofluorescence staining of 5-µm cryosections prepared from murine skin. After blocking with Roti-Immunoblock (Roth), tissue-bound rabbit IgG was visualized using Alexa Fluor 488-conjugated goat anti-rabbit IgG antibody (Invitrogen). Complement C3 deposition was detected by indirect immunofluorescence using rat anti-mouse C3 antibody (Cedarlane), followed by Alexa Fluor 546-conjugated goat anti-rat IgG antibody (Invitrogen). Coverslips were mounted with Gold mounting medium (Thermo Fisher), and fluorescence images were acquired by confocal microscopy (Leica SP5).

### Quantification of mRNA expression in skin

Gene expression in lesional skin was analyzed by RNA isolation, reverse transcription, and quantitative real-time PCR as described previously. Briefly, total RNA was isolated from skin biopsies using the RNeasy Mini Kit (Qiagen) according to the manufacturer’s instructions. After reverse transcription, cDNA was amplified on a LightCycler 480 system (Roche). For each cDNA sample and standard, 1 µL template was mixed with 9 µL master mix containing 50% 2× LC480 Probes Master, 6.25 µM sense and antisense primers, and 1% Universal ProbeLibrary probe in ddH_2_O. Primer and probe sets were selected for the respective target genes, including *Il17a*, *Il17f*, *Il1b*, *ll6*, *Cxcl1*, *Cxcl2*, and *Ccl2*. The following primer and probe sets were employed. *Hpr*t: sense 5′-TCC TCC TCA GAC CGC TTT T-3′, antisense 5′-CCT GGT TCA TCA TCG CTA ATC-3′, probe #95; *Il17a*: sense 5′-TGT GAA GGT CAA CCT CAA AGT CT-3′, antisense 5′-GAG GGA TAT CTA TCA GGG TCT TCA T-3′, probe #50; *Il17f*: sense 5′-CCC AGG AAG ACA TAC TTA GAA GAA A-3′, antisense 5′-CAA CAG TAG CAA AGA CTT GAC CA-3′, probe #463; *Il1b*: sense 5’-TGT-AAT-GAA-AGA-CGG-CAC-ACC-3’, antisense 5’-TCT-TCT-TTG-GGT-ATT-GCT-TGG-3’, probe #78; *Il6*: sense 5 ′ -GTC ACC AAA CTG GAT ATA ATC AGG A-3 ′ , antisense 5′-CCA GGT AGC TAT GGT ACT CCA GAA-3′, probe #6; *Cxcl2*: sense 5’-CCA-GCC-ACA-CTT-CAG-CCT-A-3’, antisense 5’-CAG-TTC-ACT-GGC-CAC-AAC-AG-3’, probe #49; *Ccl2*: sense 5’-CAT-CCA-CGT-GTT-GGC-TCA-3’, antisense 55’-GAT-CAT-CTT-GCT-GGT-GAA-TGA-GT-3’, probe #62; Data were analyzed using LightCycler software LCS480 version 1.5.0.39 (Roche).

### Intracellular staining of IL-17A

Skin samples were digested with Liberase TL (38 µL of 25 mg/mL; Roche) to obtain single-cell suspensions. For intracellular IL-17A staining, cells were stimulated with PMA (1:1,000), ionomycin (1:1,000), and GolgiPlug™ (1 µL; BD) for 4.5 h at 37 °C and 5% CO_2_. Cells were then stained with surface antibodies against CD45-Pacific Blue, CD4-PE-Cy7, Ly6G-BV711, γδTCR-PerCP-Cy5.5, and B220-PE (all from BioLegend), as well as CD3-FITC (BD), for 30 min at 4 °C in the dark. Subsequently, cells were fixed, permeabilized, and stained with anti-IL-17A-Alexa Fluor 647 antibody (BioLegend). Data were acquired on an LSR II flow cytometer (BD) and analyzed using FACS Express version 5 (De Novo Software).

### Stimulation of γδ T cells

γδ T cells were isolated from spleens of γδTCR-eGFP mice by fluorescence-activated cell sorting. Spleen cells were incubated with PE-conjugated antibodies against CD4, CD8, CD11b, B220, and NK1.1 (6 µL each in 200 µL; all from BD) for 30 min at 4 °C in the dark. Propidium iodide (3 µL; Miltenyi Biotec) was added immediately before sorting to exclude dead cells. γδ T cells were sorted using a FACSAria cell sorter, and data were analyzed using FACS Express version 5 (De Novo Software).

To mimic IgG and complement deposition in EBA, wells were pre-coated with murine type VII collagen (mCOL7) antigen (2 µg/well), anti-mCOL7 antibody (200 µg/well), and C3a (20 ng/well). As established inducers of IL-17A, IL-23 (20 ng/mL) and IL-1β (20 ng/mL) were added either alone or in combination to sorted γδ T cells for 48 or 72 h. IL-17A concentrations in culture supernatants were quantified by ELISA (R&D Systems).

### Statistical analysis

Statistical analyses were performed using GraphPad Prism version 6.0 (GraphPad Software). Unless stated otherwise, data are presented as mean ± standard deviation (SD). Comparisons between two groups were performed using Student’s t-test or the Mann–Whitney rank-sum test, as appropriate. For comparisons among more than two groups, one-way ANOVA followed by Bonferroni’s multiple-comparison test was used for normally distributed data. For non-parametric data, Kruskal–Wallis ANOVA on ranks followed by Bonferroni’s multiple-comparison test was applied. Statistical significance was defined as follows: *p < 0.05, **p < 0.01, and ***p < 0.001.

## ETHICS STATEMANT

All animal experiments were reviewed and approved by the Animal Research Ethics Board of the Ministry of Agriculture, Rural Areas, European Affairs and Consumer Protection of the State of Schleswig-Holstein, Kiel, Germany (approval numbers 91-9/10, 4-1/13, 12-2/14, and 109-10/19).

## DATA AVAILABILITY

The data supporting the findings of this study are available from the corresponding author upon reasonable request.

## ACKNOWLEDGMENTS

We are thankful to Nico Ghilardi (Genentech, CA) for providing IL-23p19^-/-^ mice, respectively. We are also grateful to Anja Walter and Marion Schuldt for supplying and cleaning the lab and to Ika Monath and Christine Keller for organizing the animal facility at the Research Center Borstel. This work was funded by the Deutsche Forschungsgemeinschaft (DFG, German Research Foundation) - Project number 179309734 - GRK 1727 “Modulation of Autoimmunity”. The funder had no role in study design, data collection and analysis, the decision to publish, or preparation of the manuscript.

## AUTHO CONTRIBUTIONS

Conceptualization: CH, XY, and FP; Supervision: CH; Investigation: FD, SJ, MJ, AH; Formal analysis: FD, SJ, MJ; Resources: IP, RL, and ES; Visualization: FD; Writing: FD; Writing – review & editing: CH, XY, FP, IP, RL and ES.

## DECLARATION OF GENERATIVE ARTIFICIAL INTELLIGENCE (AI) OR LARGE LANGUAGE MODELS (LLMS)

During the preparation of this manuscript, the authors used Microsoft Copilot to assist with language editing and improving readability. All AI-assisted content was critically reviewed and edited by the authors, who take full responsibility for the content of the publication.

